# RNA-seq reveals few differences in resistant and susceptible responses of barley to infection by the spot blotch pathogen *Bipolaris sorokiniana*

**DOI:** 10.1101/384529

**Authors:** Matthew Haas, Martin Mascher, Claudia Castell-Miller, Brian J. Steffenson

## Abstract

Spot blotch, caused by *Bipolaris sorokiniana* (Sacc.) Shoem., is an economically important disease affecting barley (*Hordeum vulgare* L.). The disease has largely been controlled in the Upper Midwest region of the USA through a suite of quantitative trait loci (QTL) termed the Midwest Six-rowed Durable Resistance Haplotype (MSDRH). These QTL have been bred into all six-rowed Midwest barley cultivars, including the widely used cultivar Morex. We identified a gamma ray- induced Morex mutant (MUT) that exhibits spot blotch susceptibility at the seedling stage. This mutant also spontaneously develops extremely large necrotic lesions in the absence of the pathogen at the adult plant stage. Spot blotch susceptibility at the seedling stage and necrotic lesion formation at the adult plant stage are highly correlated. To start dissecting the molecular responses underlying the observed symptoms at the seedling stage, we conducted a time course RNA-seq experiment comparing the wild type (WT) and the mutant (MUT) Morex at 12, 24 and 36 h after *B. sorokiniana* inoculation. Mock-inoculated controls were also included. A total of 10,772 and 11,530 genes were differentially expressed between treatments for WT and MUT genotypes, respectively, while 277 and 195 genes were differentially expressed between fungal and mock-inoculated genotypes, respectively. The transcript expression profiles of WT and MUT Morex samples were similar for most treatments. Two genes whose expression was putatively knocked out in the MUT were identified: HORVU3Hr1G019920 (glycine-rich protein) and HORVU5Hr1G120850 (Long- chain-fatty-acid—CoA ligase 1). The latter appears to be genetically intact, but not expressed. Collectively, these data suggest that MUT susceptibility to *B. sorokiniana* is a result of minor, rather than major, differences in the defense responses.

## Introduction

Spot blotch, caused by *Bipolaris sorokiniana* (Sacc.) Shoem., is an important disease of barley (*Hordeum vulgare* L.) and is responsible for up to 30% grain yield loss in susceptible cultivars (Mathre, 1997). However, over the past 50 years, the disease has been of little economic impact in the Upper Midwest region of the USA because widely grown six-rowed cultivars carry durable resistance (Johnson, 1984), contributed by the breeding line NDB112. Quantitative trait loci (QTL) mapping and genome-wide association studies (GWAS) have revealed that the NDB112-derived resistance is conferred by three QTL on chromosomes 1H, 3H and 7H, collectively referred to as the Midwest Six-rowed Durable Resistance Haplotype (MSDRH) (Steffenson et al., 1996, Bilgic et al., 2005, Roy et al., 2010, Zhou and Steffenson, 2013). The spot blotch resistance gene *Rcs5* confers seedling resistance to *B. sorokiniana* isolate ND85F (Steffenson et al., 1996) and co-locates with the chromosome 7H QTL (*Rcs-qtl-7H-11_20162*) of the MSDRH (Zhou and Steffenson, 2013). High resolution mapping of the *Rcs5* region by Drader (2010) revealed two wall- associated kinases, suggesting a putative function of *Rcs5.* Efforts to clone and functionally validate *Rcs5* are ongoing (R. Brueggeman, personal communication).

Despite the wealth of knowledge about the genetic architecture of durable spot blotch resistance, little is known about the underlying molecular basis of resistance to spot blotch (Drader, 2010). Another study that examined defense mechanisms against spot blotch was done using the microarray Affymetrix Barley1 GeneChip hybridized with seedling transcripts of the resistant wild barley (*Hordeum vulgare* ssp. *spontaneum*) accession Shechem 12-32 (Millett et al., 2009). That study included three early time points of pathogen colonization starting at 12h to 36h and found, within the defense category of the GO annotation, PR proteins (i.e. chitinases), receptor-like kinases, PAL related, proteinase inhibitor, resistance (R) genes and defense related transcripts accumulated at different time points in the pathogen-infected treatment compared to the mock-inoculated (water) control. No transcripts mapped to the known QTL (*Rcs-qtl-7H-11_20162*) for seedling resistance in 7H chromosome.

RNA-seq technology enhances the likelihood of finding more transcripts at different levels of expression; thus, transcriptome profiling can be used to elucidate the expression of genes associated with this valuable source of spot blotch resistance in barley at the seedling stage. RNA-seq has since supplanted microarrays and has been applied in many different pathosystems including soybean-bacterial leaf pustule (Kim et al., 2011), cotton-Verticillium wilt (Xu et al., 2011), and potato-late blight (Gao et al., 2013). Each of these studies differs slightly in their experimental design. For example, Kim et al. (2011) compared near-isogenic soybean lines (NILs) and Gao et al. (2013) compared ‘Russet Burbank’ potatoes with and without the transgene *+RB*. Xu et al. (2011), on the other hand, used completely different species of cotton: the resistant *Gossypium barbadense* and the susceptible *G. hirsutum*. Each study revealed transcript expression patterns specific to the individual pathosystem, but all found that resistant plants were able to mount a more rapid and sustained defense response against the invading pathogens.

To further understand the molecular bases behind the Morex barley–*B. sorokiniana* pathosystem, we initiated a study that involved two phases: identification of spot blotch susceptibility mutants and then characterization of the mutants by RNA-seq. We searched for spot blotch susceptibility mutants in a gamma ray-irradiated population of Morex barley grown in a field nursery inoculated with *B. sorokiniana* isolate ND85F. Several putative susceptibility mutants at the adult stage were identified in the field and then tested in the greenhouse at the seedling stage to corroborate the reaction. One mutant, which we refer to as Morex Mutant 14-40 in our lab (MUT), in particular showed clear spot blotch susceptibility at the seedling stage after challenge with *B. sorokiniana* isolate ND85F. As this MUT was being grown for seed, it developed very large spontaneous lesions that mimicked those caused by the pathogen (i.e. oval necrotic areas surrounded by chlorosis that later coalesced). A selected mutant progeny exhibiting clear spot blotch susceptibility at the seedling stage and large spontaneous lesions at the adult plant stage was subjected to RNA-seq profiling to identify transcripts that were differentially expressed compared to those of the Morex wild type (WT). The objective of this study was to identify genes with altered expression patterns between both genotypes WT and MUT at the seedling stage that could increase our understanding of the underlying basis of resistance to spot blotch.

## Methods

### Plant material and experimental setup

The spot blotch resistant barley genotype used in these experiments was the six-rowed malting cultivar ‘Morex’ (Rasmusson and Wilcoxson 1979), which derives its resistance from the breeding line NDB112 (CIho 11531). NDB112 originated from the cross CIho 7117-77/Kindred (Wilcoxson et al., 1990) and is highly resistant to most pathotypes of *B. sorokiniana* (Valjavec-Gratian and Steffenson 1997). The WT Morex used in these experiments was from the original ‘Morex’ seed source used to create the Steptoe/Morex doubled haploid (DH) population (Kleinhofs et al., 1993). To generate mutants susceptible to spot blotch, 3,000 WT seeds were irradiated with 150 gray gamma radiation by A. Kleinhofs at Washington State University in 2005, and then the M_1_ generation was increased in the field at Pullman, WA during the same year. The resulting M_2_ families were sown at the Minnesota Agricultural Experiment Station (MAES) on the Saint Paul campus of the University of Minnesota in 2008. The M_2_ nursery was inoculated with *B. sorokiniana* (isolate ND85F) according to established protocols (Steffenson et al., 1996) to identify families segregating for reaction to the disease. A few families segregating for spot blotch severity and infection response (IR) were identified in the nursery. One line within a segregating family was particularly interesting due to its high spot blotch severity that suggested susceptibility at the adult stage in the field. Cleaved amplified polymorphic sequences (CAPS) markers, specific for Morex, were used to initially confirm the WT genetic background of the mutant. Subsequent crossing back (one generation, BC_1_) to the Morex was performed in order to increase homozygosity and standardize the genetic background of the MUT with respect to non-target mutations, but these BC progenies were not subjected to RNA- seq. In doing so, we did not recover any family that with the same high level of resistance as WT Morex after *B. sorokiniana* infection at the seedling stage. Highly susceptible families, however, were readily recovered. Through observations of the development of these plants in the greenhouse, we discovered that the MUT plants developed extremely large necrotic lesions at the adult plant stage very similar to spot blotch symptoms (Figure 1). The plants exhibiting these large spontaneous lesions at the adult plant stage also were highly susceptible to spot blotch at the seedling stage, indicating an association between the two traits. The genomic locus/loci of the mutations in the MUT are not known at this time.

**Figure 1.**
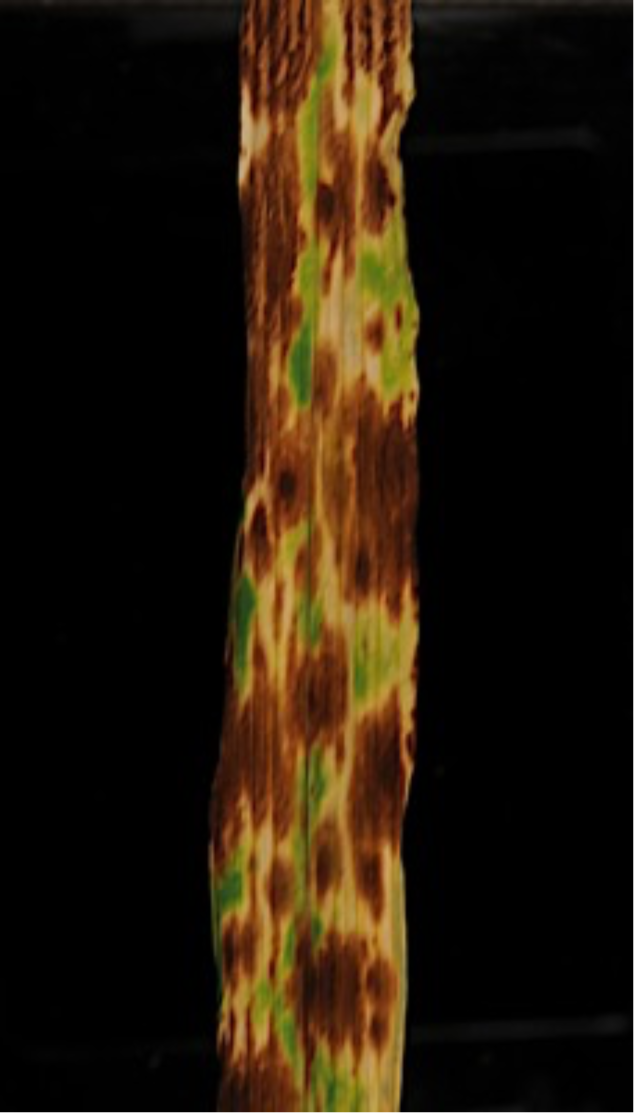
Necrotic lesions of the adult Morex MUT 14-40 grown in the greenhouse.

One experimental unit consisted of fifty individual plants sown individually in plastic cones (5 cm diameter × 18 cm depth, Stuewe and Sons, Inc., Tangent, OR, USA) filled with a 50:50 mixture of steam-sterilized soil and Sunshine MVP potting mix (Sun Gro Horticulture, Quincy, MI, USA). Forty of these individuals were either WT or MUT and used in the transcriptome study. The remaining plants were used to verify the disease reaction after inoculation with *B. sorokiniana* isolate ND85F, but were not harvested for transcriptomic analysis. These plants included controls NDB112 (CIho 11531) and Bowman (PI 483237), both resistant to ND85F; ND5883 (PI 643237), susceptible to the isolate (Fetch and Steffenson, 1999) and the WT and MUT genotypes. Plants were organized in cone racks in a randomized complete block design (RCBD). Plants were grown for 14 d until the second leaf of each seedling was fully expanded.

Inoculum was prepared according to the methods of Fetch and Steffenson (1999). Briefly, conidia of *B. sorokiniana* isolate ND85F, retrieved from silica gel crystals stored at 4°C, were plated out onto minimal media agar plates 10 d prior to inoculation. To harvest conidia for inoculation, water was poured onto the surface of the *B. sorokiniana* cultures and then a sterile rubber spatula was used to dislodge conidia. This conidia-water suspension was then passed through four layers of cheesecloth to remove mycelial fragments and the concentration estimated using a hemacytometer. For inoculation, the suspension was diluted to 6,000 conidia/ml and a surfactant (Tween-20^®^: polyoxyethylene-20-sorbitan monolaurate at 10 μl / 100 ml of suspension) added to increase the distribution and adhesion of conidia to the leaf surfaces. Approximately 0.3 ml of the conidial suspension was applied to each inoculated leaf.

At the time of inoculation, the mock-inoculated rack was treated first, immediately followed by the inoculated rack. This was done to avoid possible cross-contamination of the mock-inoculated set. After inoculation, the plants were moved to a mist chamber kept near 100% relative humidity by ultrasonic humidifiers running for 2 min of continuous misting every 60 min. During this period, the plants were kept in the dark overnight (16 h). After the mist treatment, the doors to the mist chambers were opened and the plants were allowed to dry slowly (4 h) before being moved back to the greenhouse. The first tissue samples were harvested 12 h after inoculation (hai) while the plants were still in the mist chamber. Subsequent samples were collected at 24 and 36 hai. The samples were immediately frozen in liquid nitrogen and then stored at - 80°C until RNA extraction was performed. In total, 36 independent samples were collected from three biological replicates of two treatments (*B. sorokiniana*-inoculated and water-inoculated), two plant genotypes (WT and MUT) and three time points (12, 24 and 36 hai). Time points were selected based on a previous study that indicated *C. sativus* switches its lifestyle from that of a biotroph to a necrotroph around 24 hai (Millett et al., 2009). The 12 and 24 hai time points were selected to capture potentially interesting defense responses at even intervals before and after the 24 hai time point.

### Sample preparation

Total RNA was extracted using a modified TRIzol method. First, tissue samples were ground into a fine powder with a mortar and pestle in liquid nitrogen. This powder was then transferred to a 50 ml tube with 10 ml TRIzol reagent (Ambion, Foster City, CA, USA) and kept on ice. Next, the samples were incubated for 15 min in a pre-heated water bath at 60°C. The tubes were then spun for 10 min at 4,000 rpm, and the supernatant transferred to a RNase-free 50 ml tube to which 2 ml of chloroform was added. The tubes were gently inverted 5-10 times, centrifuged for 15 min at 4,000 rpm, and the resulting supernatant transferred to a new RNase-free 50 ml tube. One half of one volume of isopropanol and Na-citrate/NaCl solutions were added per 1 ml of aqueous solution followed by incubation at -20°C for 20 min. Then, samples were centrifuged for 10 min at 4,000 rpm, rinsed with 10 ml of 70% EtOH and centrifuged for another 5 min at 4,000 rpm. The supernatant was then discarded and the pellet was allowed to air dry for at least 10 min while inverted on a paper tissue. Finally, the pellet was dissolved in 300 µl DEPC-treated (RNase-free) water and frozen at -20°C until all samples were ready for sequencing. Upon thawing, 2 µl RNasin (Promega, Madison, WI, USA) was added to each sample. RNA was purified using an Rneasy Cleanup Kit (Qiagen, Foster City, CA, USA).

RNA purity was determined using a NanoDrop™ 2000 spectrophotometer (Thermo Fisher Scientific, Inc., Wilmington, DE, USA), and the integrity in a 2100 Agilent Bioanalyzer™ (Agilent Technologies, Santa Clara, CA, USA). Preparation of cDNA libraries was conducted using TruSeq RNA v2 library kits. Libraries were gel size selected to have insert sizes of 200 bp. Thirty-six samples were sequenced (100 bp paired-end) on an Illumina HiSeq 2000, and one sample (R3_INOC_MUT_24) was re-sequenced on an Illumina HiSeq 2500 by the University of Minnesota Genomics Center (UMGC).

### Read alignment and sequence analysis

The BAC-based barley reference sequence (Mascher et al., 2017) was used to align RNA-seq data. Read count and TPM (transcript per million reads mapped) data were determined using Kallisto software version 0.43.0 (http://pachterlab.github.io/kallisto) (Bray et al., 2016), and the resulting abundance data were imported into the R statistical environment version 3.3.2 (R Core Team, 2016) for statistical analysis. Data were normalized using edgeR (Robinson et al., 2010) and limma (Ritchie et al., 2015). Genes with less than 100 counts across all samples were excluded from further analysis. The voom (variance modeling at the observational level) transformation was used to account for the mean-variance relationship of the count data. One thousand genes with the highest variances were used to calculate the row variance through Principal component analysis implemented in the R package matrixStats (Bengtsson 2017). Differential expression analyses were conducted by fitting the voom-transformed count matrix to the linear model specified for the design matrix and performing gene contrasts of interest. The Benjamini-Hochberg method was used for error correction due to multiple testing (Benjamini and Hochberg, 1995). For each time point, differential gene expression (i.e. between genes for WT vs, MUT genotypes, and between INOC vs. MOCK of the same genotype) were calculated. Genes were determined to be differentially expressed when the log_2_ fold change was greater than 2 or less than -2 with an adjusted *p*-value of less than 0.05. All raw sequence data are available from the National Center for Biotechnology Information Short Read Archive (NCBI SRA) under accession number SRP071745.

### Validation of gene expression with qPCR

We selected 10 genes to validate the differential expression results. Three independent experiments were conducted according to the same methods used for the original RNA-seq experiments. After total RNA extractions from the second leaves of both WT and MUT plants, conversion to cDNA was done using the Superscript first-strand synthesis system (Invitrogen, Foster City, CA, USA). Primers were designed using the NCBI primer design tool (Ye et al., 2012). The RNA-seq data were initially analyzed using the previous whole-genome shotgun (WGS) barley reference sequence (International Barley Genome Sequencing Consortium, 2012), and therefore the primers for qPCR were designed using the MLOC designators, rather than the HORVU designators from the updated BAC-based barley reference genome, which became available during the course of this project. The qPCR results were carried out at the UMGC using the Roche Universal Probe Library (UPL) system, which is functionally similar to the TAQMAN assay. The Roche UPL system is more accurate than SYBR Green because each probe (8-9 nt) consists of two parts: a reporter (fluorescein) at the 5’ end and a quencher dye at the 3’ end of the probe. Upon integration of the probe into the transcript during qPCR, the quencher is cleaved as a result of integration into a double-stranded cDNA molecule. The intensity of the signal is therefore related to the number of integrated probes. Analyses were performed on an ABI 7900HT machine (Applied Biosystems, Foster City, CA, USA). Raw C_T_ values were used to determine log_2_ of fold-change values according to the method of Schmittgen and Livak (2008). Only two replicates were available for the qPCR. because of problems with the RNA quality of the third replicate. The actin and glyceraldehyde-3-phosphate dehydrogenase genes were used as housekeeping genes for normalization. Primer information for genes used in qPCR validation is provided in Table 1.

**Table 1.**
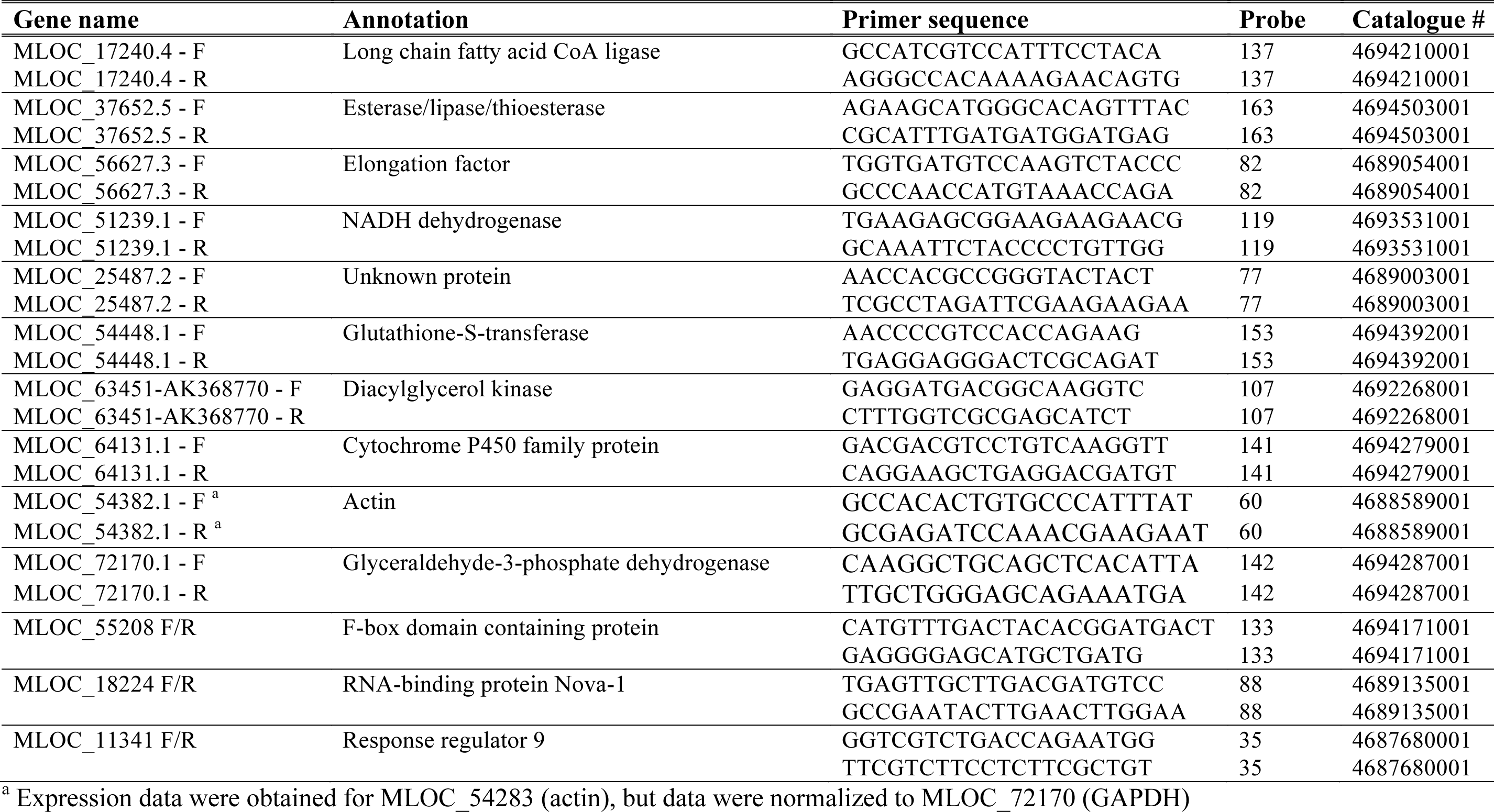
Primer information for qPCR validation of selected genes using the Roche Universal Probe Library (UPL) system at the University of Minnesota Genomics Center.

### Identification of putative knockout

Based on transcriptome analyses, two genes HORVU3Hr1G019920 (glycine-rich protein) and HORVU5Hr1G120850 (Long-chain-fatty-acid—CoA ligase 1) were identified as putative knockouts because their expression was abolished in the MUT but not in the WT across both treatments (Figure 2). Primers were designed to test for the presence of HORVU5Hr1G120850 (MLOC_17240) (Table S1) based on the 2012 WGS reference sequence (IBGSC, 2012). The MUT was not tested for the presence of HORVU3Hr1G019920 because its annotation was not informative (i.e., a glycine rich repeat). DNA extracted from WT and MUT was sent to Functional Biosciences, Inc. (Madison, WI, USA) for PCR and sequencing of products according to their methods. Quality trimmed FASTA files were aligned to the complete sequence of MLOC_17240 using Clustal Omega (Supplemental File S1). The sequenced products were deposited in the NCBI SRA under reference numbers TI 2344113413-2344113439.

**Figure 2.**
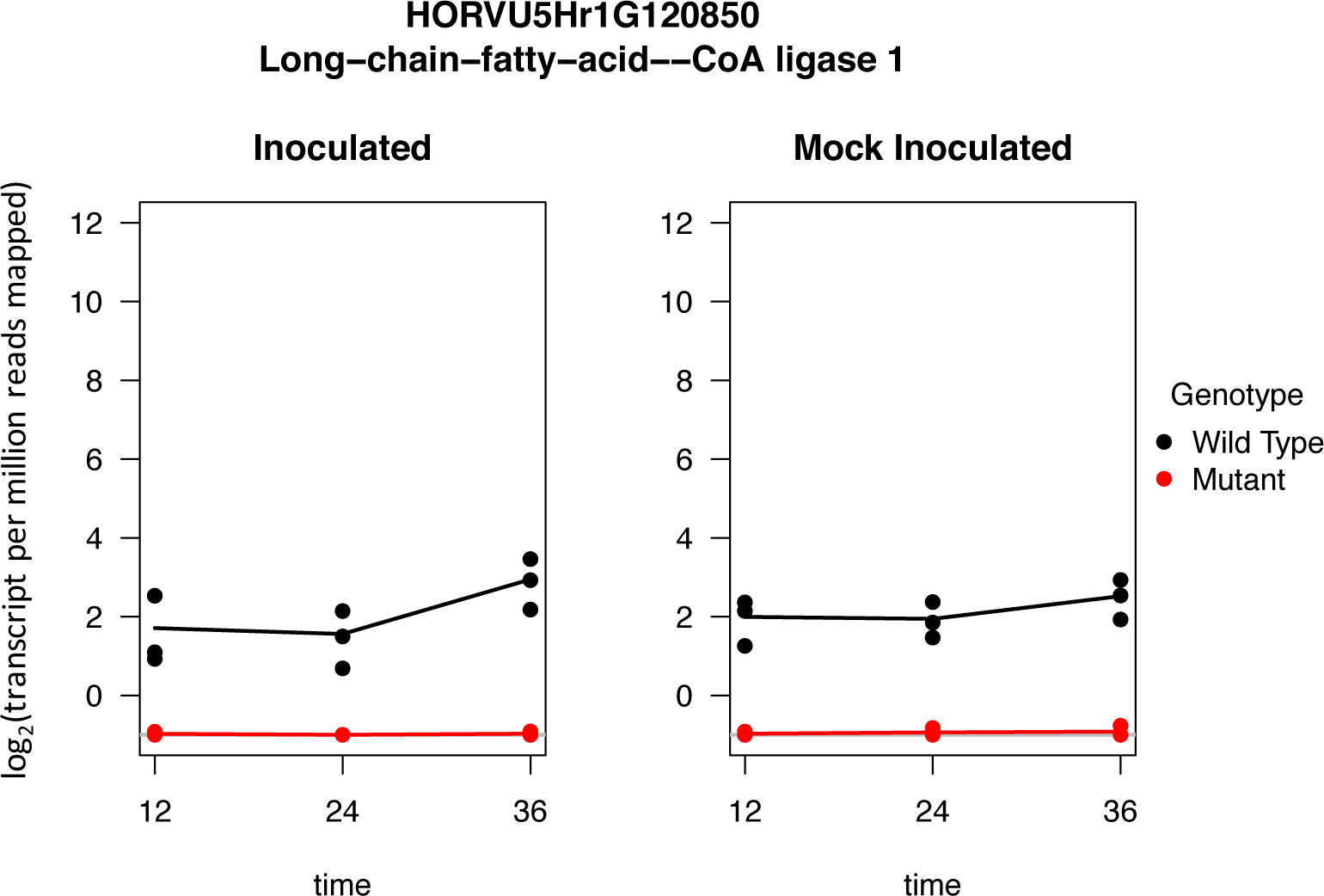
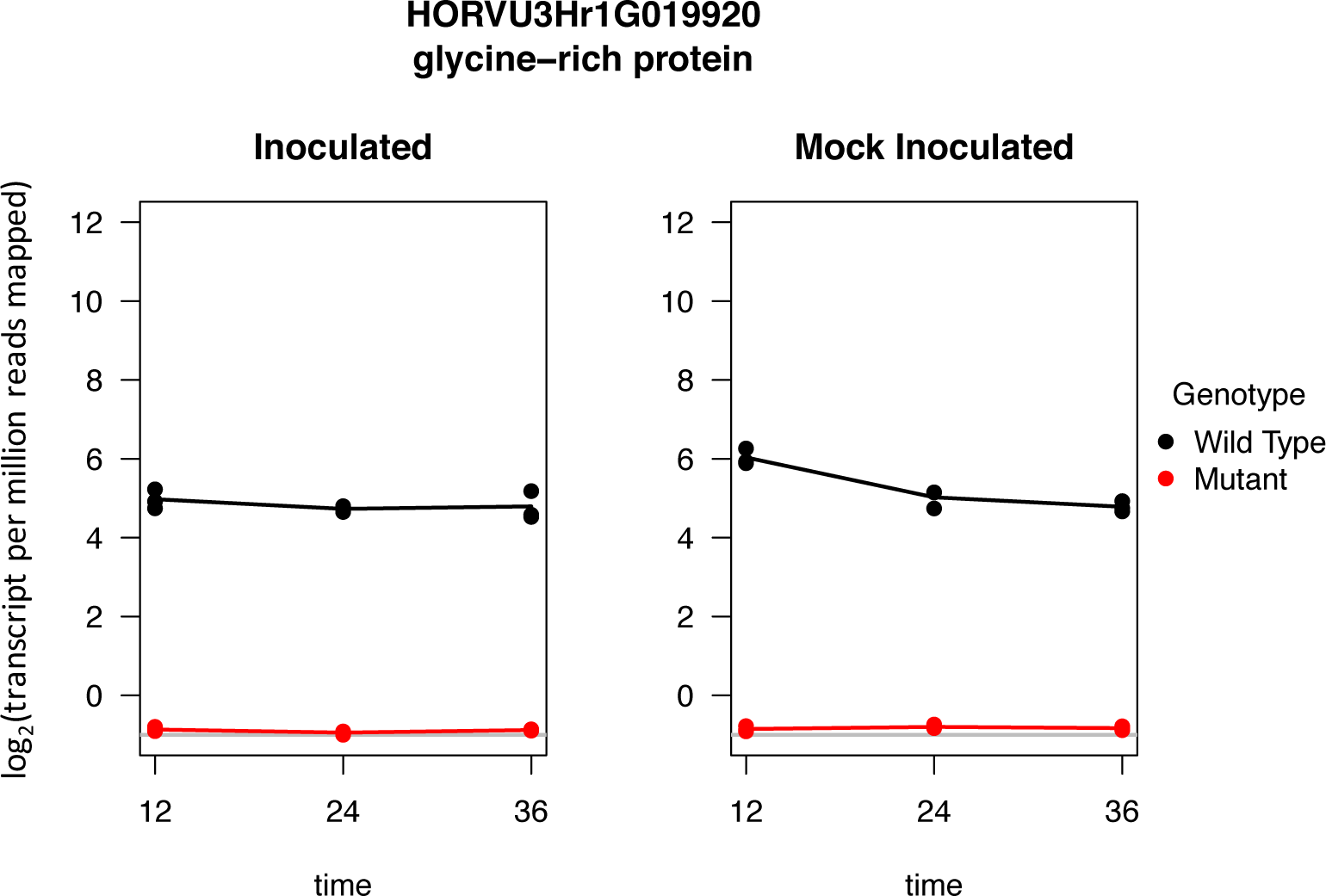
Expression profile of A) HORVU5Hr1G120850 (top) and B) HORVU3Hr1G019920 (bottom) for the WT and MUT genotypes at three time points: 12, 24 and 36 hours after inoculation (hai) and mock-inoculation. The plots show abolished expression of both genes in the MUT for both treatments at all time points.

## Results

### Symptom development

Disease symptoms developed as expected on the inoculated 14-day old WT and MUT seedlings that were not collected for RNA extraction. WT plants showed an infection response (IR) of 3 with a minimal amount of associated chlorosis (Figure 3A), while MUT plants had an IR of 6-7 with a high degree of chlorosis (Figure 3B). The controls of NDB112 (IR = 2-3), Bowman (IR = 4-5) and ND5883 (IR = 7-8) also reacted as expected, indicating that the inoculation with *B. sorokiniana* isolate ND85F was successful. The water-treated (mock-inoculation) plants did not develop any disease symptoms after 12 days, as expected (Figure S1).

**Figure 3.**
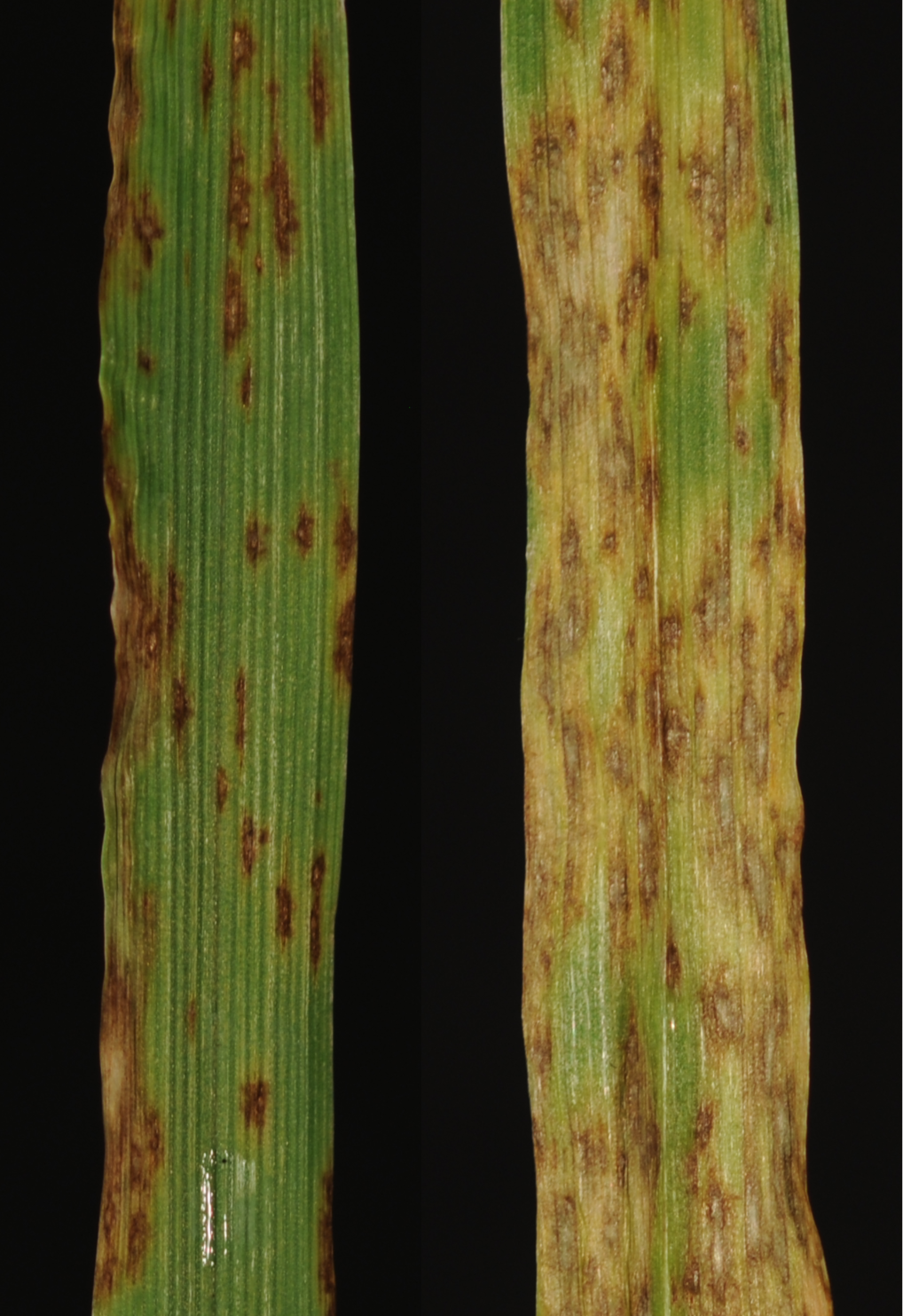
The second leaves of A) WT Morex (left) and B) MUT Morex (right) responses to *C. sativus* isolate ND85F 12 days after inoculation. The reaction of the WT is indicative of a resistant reaction, while the reaction of the MUT is indicative of a susceptible reaction.

### Principal component analysis

The first principal component (PC) explained 69% of the variance and separated the inoculated and mock-inoculated samples (Figure 4, left). The second principal component explained 10% of the variance and separates the 12-hour inoculation time point from the 24- and 36-hour time points (Figure 4, right). The third and fourth principal components explained a combined 7% of the variance and separated the samples into four clusters: 1) Mock-inoculated WT and MUT genotypes at the 24-hour time point (Figure 5), 2) Inoculated and mock-inoculated samples at the 12- and 24- hour time points, 3) Inoculated and mock-inoculated samples of both genotypes at the 12- and 36-hour time points and 4) Inoculated WT and MUT genotypes at the 12- and 36-hour time points (Figure 5).

**Figure 4.**
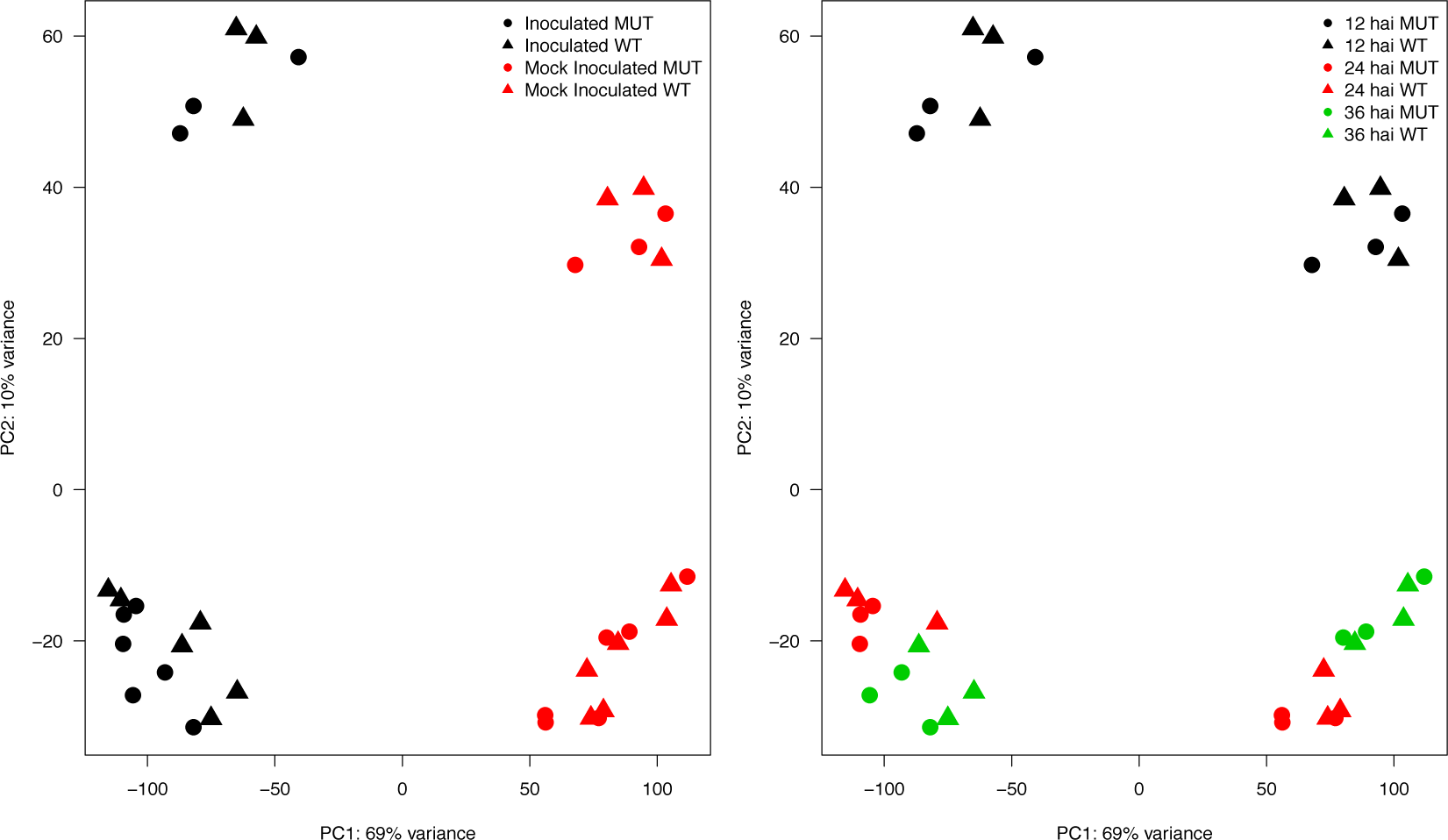
Principal components 1 and 2 show that the samples cluster according to treatment (left) and by time after inoculation (right). The 12-hour time point is most different from the 24- and 36-hour time points. WT and MUT genotypes from each time point cluster together.

**Figure 5.**
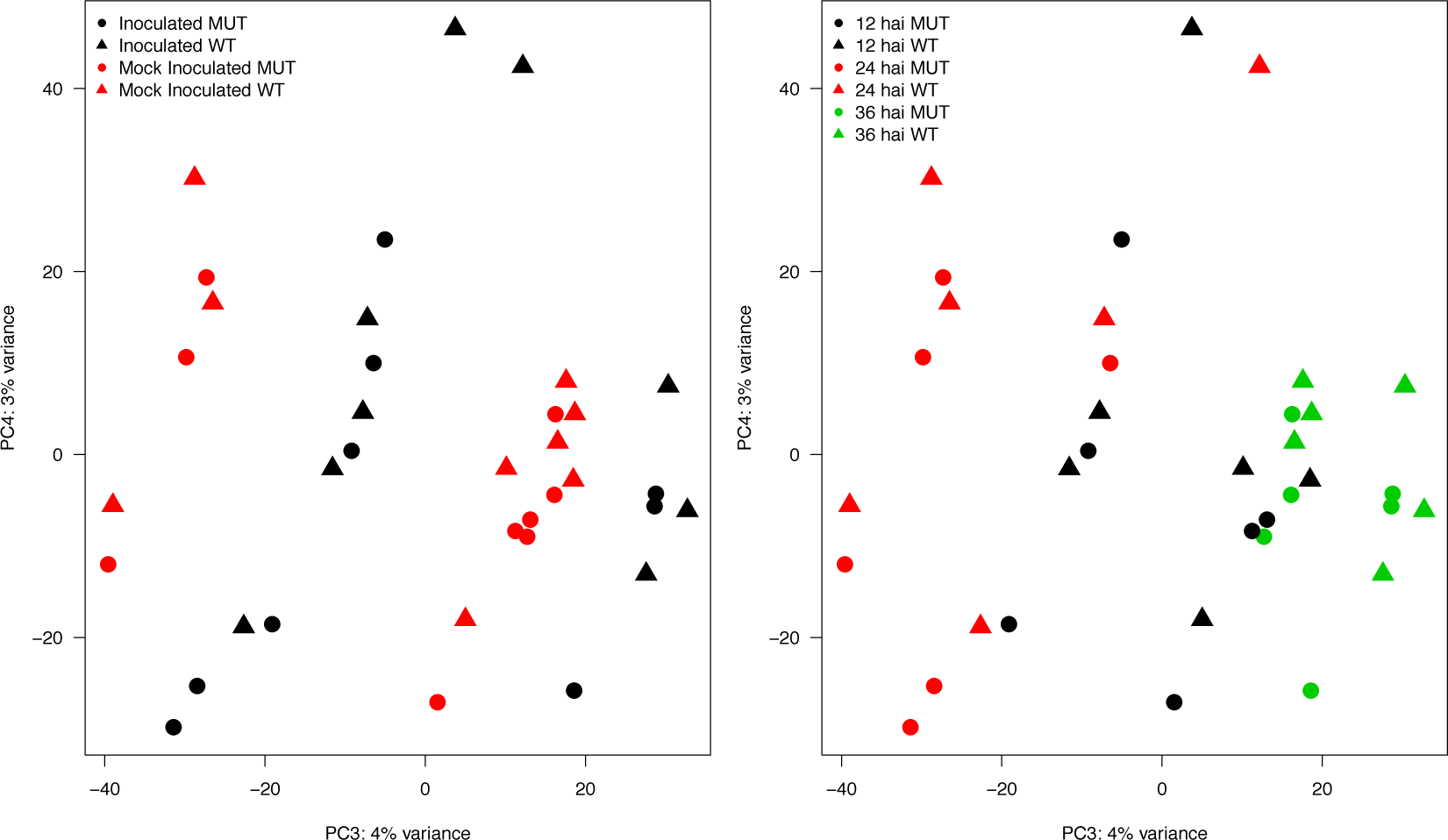
Principal components 3 and 4 explain a small amount of the variance (4% and 3%, respectively), but they also show that samples cluster by treatment (left) and time after inoculation (right).

### Differential expression between genotypes

A total of 325 genes were differentially expressed between genotypes among INOC samples, including 78, 147 and 100 genes at 12, 24 and 36 hai, respectively (Figure S2A). Of these 325 genes, 44 were differentially expressed at all time points evaluated, while 17, 147, and 19 were differentially expressed exclusively at 12, 24, and 36 hai. A total of 345 genes were differentially expressed between genotypes among MOCK samples, including 102, 140 and 103 genes at 12, 24 and 36 hai, respectively (Figure S2B). Two genes were of particular interest from this analysis: a glycine-rich protein (HORVU3Hr1G019920) and a long chain fatty acid CoA ligase (HORVU5Hr1G120850). HORVU3Hr1G019920 and HORVU5Hr1G120850 were among the highest differentially expressed genes due to their expression being completely abolished in the MUT. However, these genes showed a similar expression pattern when mock-inoculated, so expression in the WT is likely not influenced by treatment with *B. sorokiniana* (Figure 2).

### Differential expression between treatments

A total of 10,772 genes were differentially expressed between treatments among WT samples, including 1,947, 1,562 and 1,283 unique genes at 12, 24 and 36 hai, respectively and 2,802 expressed at all three time points (Figure S3C). A total of 11,530 genes were differentially expressed between treatments among MUT samples including 2,059, 1,143 and 1,897 unique genes at 12, 24 and 36 hai, respectively and 3,415 expressed at all time point (Figure S3D).

### Validation with qPCR

Eight genes were selected for validation by qPCR because they exhibited the highest level of differential expression using RNA-seq data. Some of the genes selected for validation by qPCR agree with the RNA-seq differential expression results (Figure 6). Several genes including an esterase/lipase/thioesterase family protein (HORVU7Hr1G018470), a diacylglycerol kinase 5 (HORVU7Hr1G092530), an NBS- type putative resistance protein (HORVU3Hr1G052120), and an elongation factor (HORVU1Hr1G003960) show no apparent difference in expression, as measured by log_2_ fold change, despite being differentially expressed in the RNA-seq samples. These differences could be the result of different statistical procedures used to calculate log_2_FC for RNA-seq data and qPCR data. One gene, a glutathione S-transferase family protein (HORVU7Hr1G002720), showed a modest down regulation in the WT compared to the MUT by RNA-seq and a steep down regulation (to a log_2_FC value of -88) at the 36 hai time point. This result may also be a statistical artifact, although the SYBR green method was used initially for some of these genes and HORVU7Hr1G002720 also showed significant down regulation at the 36 hai time point (*data not shown*). For a long chain fatty acid CoA ligase (HORVU5Hr1G120850/MLOC_17240), no transcript could be amplified from the mutant samples. Due to the absence of this transcript, no CT values could be determined, resulting in our inability to compute log_2_FC values for these samples. Log_2_FC values were able to be calculated for RNA-seq data only because a value of 0.5 was added to individual gene counts from all samples prior to analysis in order to account for genes with a count of 0. Something similar could be done with the qPCR data (such as imputing a CT value of ~45 in place of ‘NA’ values) but since this transformation would not be applied to all samples, it would insert bias into the data. Two genes, a Sec14p-like phosphatidylinositol transfer family protein (MLOC_34706) and a disease resistance protein (MLOC_68128, on chromosome 1H) could not be validated because even despite several (6) attempts, a suitable probe could not be designed to obtain meaningful data. Probes were deemed unsuitable because the PCR efficiency (based on dilution tests) were too high to be reliable.

**Figure 6.**
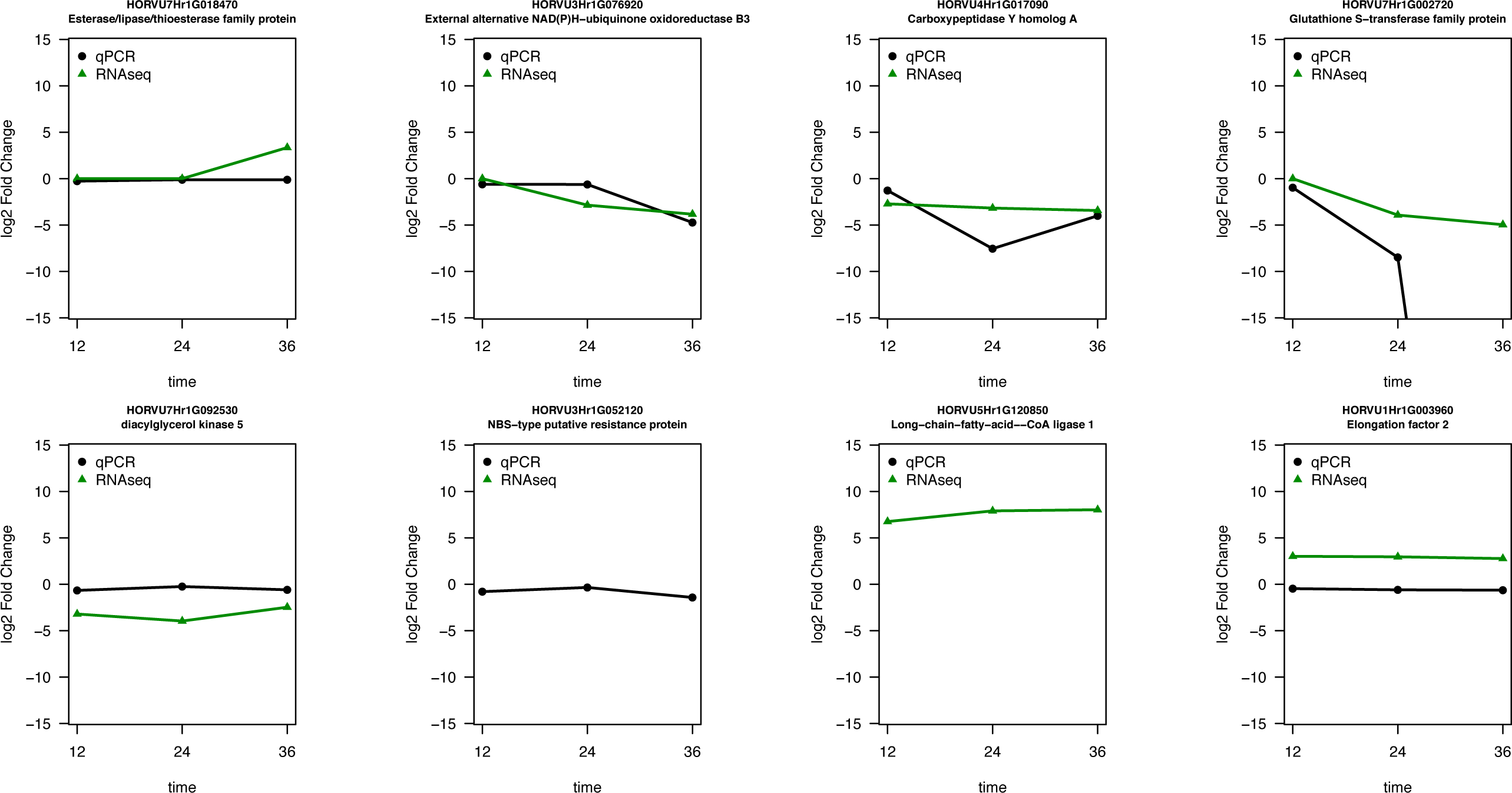
Log_2_ fold-change plots for select genes showing values obtained through qPCR (black circles) and RNA-seq (green triangles).

### Identification of putative knockout

Four of the six primers designed to test for the presence of HORVU5Hr1G120850 (MLOC_17240) in the MUT amplified products from MUT genomic DNA, but some primers did not amplify in the mutant (17R, 16F and 16R). Multiple genes with the same annotation exist, including two others (HORVU5Hr1G040740 and HORVU5Hr1G046950) on chromosome 5H; however, the sequences are not similar enough that the PCR product could have originated from one of them rather than from HORVU5Hr1G120850.

## Discussion

The spot blotch resistance of six-rowed barley cultivars grown in the Upper Midwest region is remarkable for its longevity and is conferred by the MSDRH. To identify genes with altered expression patterns that could increase our understanding of the underlying basis of resistance to spot blotch, we used a unique Morex MUT to study transcriptional changes in comparison with the WT at the seedling stage during the early infection period by *B. sorokiniana* as well as the simultaneously mock-inoculated plants. Several genes of interest, including the long-chain fatty acid CoA ligase (MLOC_17240/HORVU5Hr1G120850) were differentially expressed between the WT and MUT genotypes during the early infection period; however, many differentially expressed genes between WT and MUT after inoculation were also differentially expressed between WT and MUT genotypes under the mock-inoculated conditions. Therefore, we cannot conclude that the differential expression of these genes is a response to *B. sorokiniana*; however, at least some of these genes may show constitutive differences throughout the development of the WT and MUT plants. Some classes of genes, such resistance genes, would be expected to be expressed at all times if they act as guards or sentinels to intercept signals from pathogenic microbes.

The number of differentially expressed genes observed for each contrast matches our expectations based on the PCA plot (Figures 4-5). The greatest number of differentially expressed genes was found from the contrasts between inoculated and mock-inoculated samples. The first PC separated these two groups. The second PC separated the 12-hai samples from the 24- and 36-hai samples. A likely explanation for this is that the 12-hai samples were collected while the plants were still in the inoculation chambers where the humidity was near 100%. Therefore, any conclusions based on differential expression at the 12-hai time point should take this into consideration. The third and fourth PCs further separated samples based on time points, but did not separate samples based on genotype. From this, we may infer that the expression profiles of the WT and MUT samples are very similar to one another. The small number of differentially expressed genes between the WT and MUT samples and the similarity of their respective expression profiles provides further evidence that the two genotypes are very similar.

The paucity of differentially expressed genes between the WT and MUT genotypes was surprising. This a few possibilities could explain these results. First, the two genotypes used in this study are nearly identical. Both genotypes are the cultivar Morex, but one has been mutagenized, resulting in susceptibility at the seedling stage when inoculated to *B. sorokiniana* isolate ND85F, when remaining resistance at that stage in the WT. In addition of that phenotypic change, the MUT develops lesions that mimic spot blotch at the adult stage while the developmental states of the MUT are similar to those of the WT. At this point the genetic changes putatively induced by mutated locus/loci are still unknown. A second possibility is that, although the goal of this work was to study the early infection response, the time points selected were too early for enough defense-related transcripts to have accumulated. Thus, most of the cells in the barley leaf would not yet be responding to inoculation.

Products from HORVU5Hr1G120850 (MLOC_17240) were amplified in the MUT, ruling out a deletion of the entire gene. Small differences between both the WT and MUT genotypes would be expected; however, the differences observed in primer pair 20F and the fact that some of the primers did not amplify products in the MUT, but did in the WT suggest that there may be a smaller deletion within the gene itself. Such a deletion could have caused a frame shift resulting in missense mutations or a truncated transcript. In either case we might still expect to detect these mutated transcripts by RNA-seq, even if the protein translated from them is non-functional. Alternate explanations include that the mutant transcript is rapidly degraded or that an alteration in the 5’-UTR prevents transcription from taking place. This MUT is an interesting genetic resource for the barley research community, both for its altered spot blotch reaction at the seedling stage, but perhaps especially for the adult-onset necrotic lesions. Future work should investigate the genomic position(s) of the mutation through genetic mapping of reaction to *B. sorokiniana* as well as the severe necrotic lesions. Whole-genome shotgun sequencing of the MUT may also be used to detect differences between the MUT and the Morex reference genome.

### Data availability

Data from this experiment are available online through e!DAL (Arend et al., 2014) at the following URL: https://doi.ipk-gatersleben.de/DOI/c0602305-58bb-4029-bd8d-148bbf381b40/8bbc5b5f-3a1d-4c55-bceb-44229361d429/2/1847940088. The directory contains two Rdata files. 170309_expression_data.Rdata contains the expression (count) matrix and sample info from the RNA-seq experiment. 170424_morex_mut_qpcr.Rdata contains raw and transformed expression data obtained using qPCR. A final DOI will be assigned to these data before final publication of this manuscript.

## Acknowledgements

The authors would like to thank Dr. Nirmala Jayaveeramuthu for excellent guidance with RNA extraction and qPCR experiments. Computing resources from the Minnesota Supercomputing Institute and especially the Leibniz Institute of Plant Genetics and Crop Plant Research (IPK) are greatly appreciated.

## Author contributions

**Conceptualization:**Brian Steffenson

**Formal analysis:**Matthew Haas, Martin Mascher

**Funding acquisition:**Brian Steffenson

**Investigation:**Claudia Castell-Miller, Matthew Haas, Brian Steffenson

**Resources:**Brian Steffenson, Martin Mascher

**Supervision:**Brian Steffenson, Martin Mascher

**Writing-Original draft preparation:**Matthew Haas

**Writing-Review and editing:**Matthew Haas, Martin Mascher, Claudia Castell-Miller, Brian Steffenson

---

**Figure S1.**
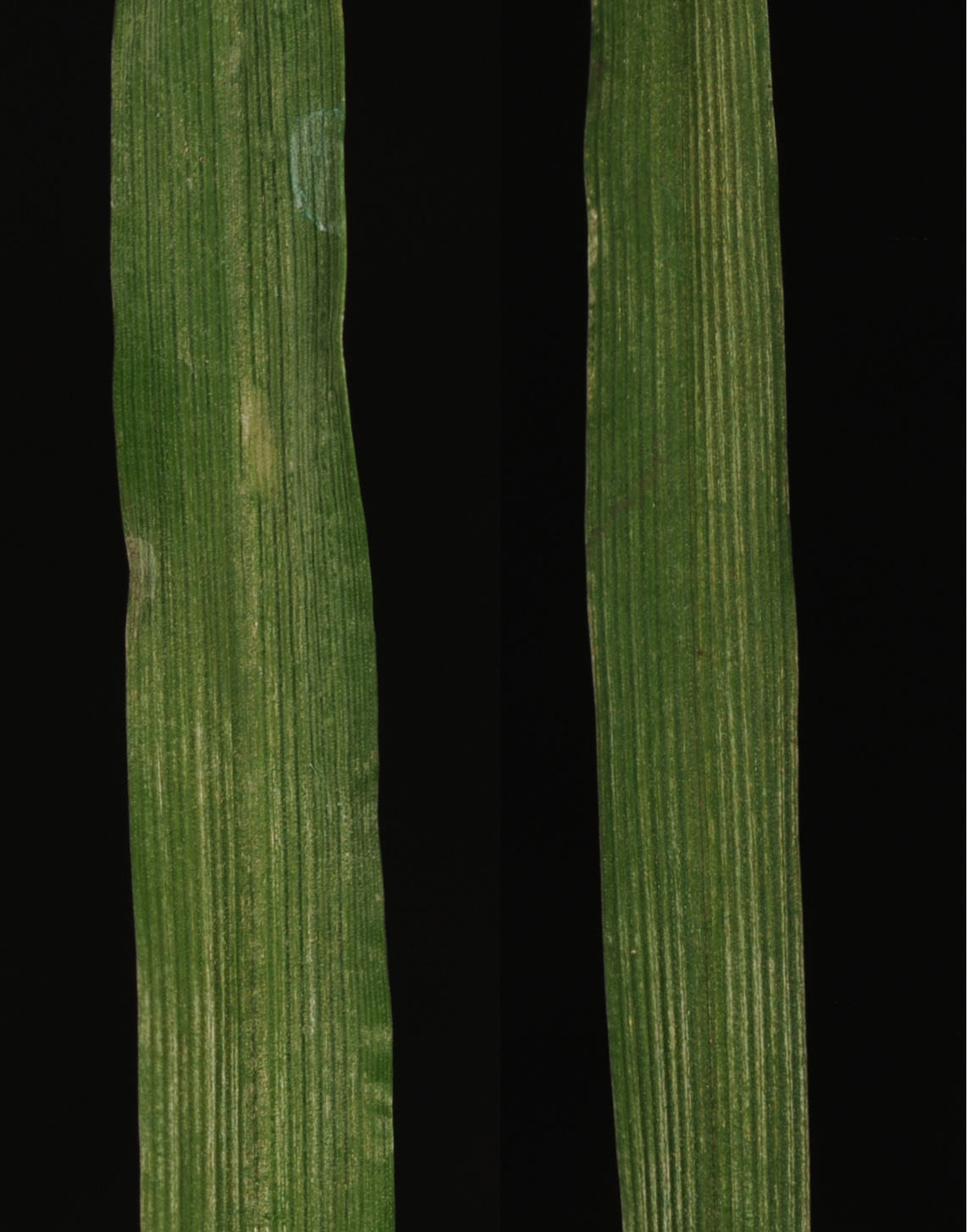
The second leaves of WT (left) and MUT (right) plants exhibited no symptoms 12 days after mock-inoculation.

**Figure S2.**
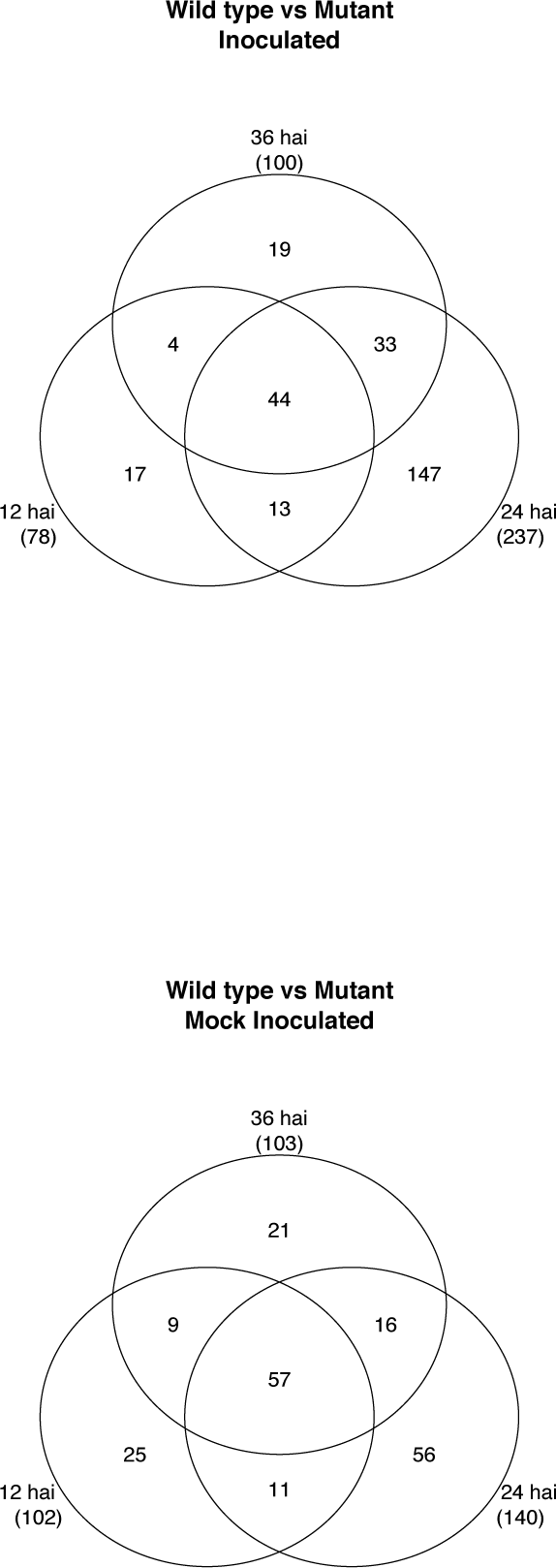
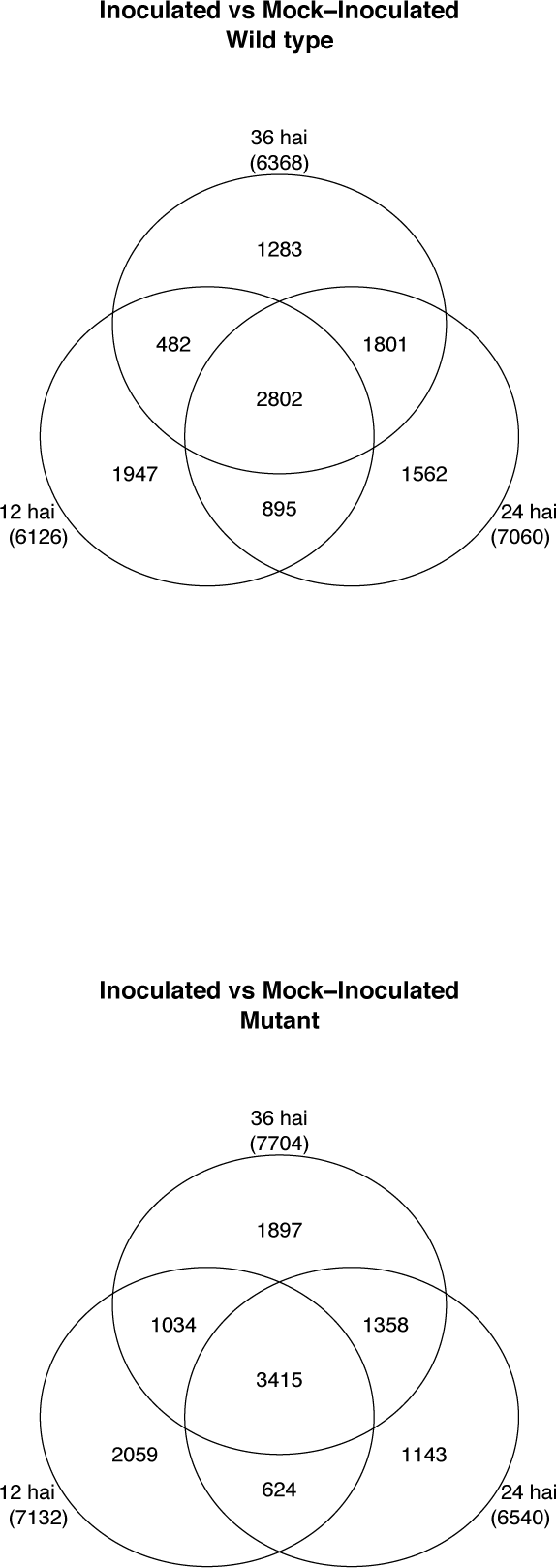
Venn diagrams for (A) Inoculated WT vs MUT samples, (B) Mock- Inoculated WT vs MUT samples, (C) WT Inoculated vs. Mock-Inoculated samples, and (D) MUT Inoculated vs. Mock-Inoculated samples.

**Table S1.**
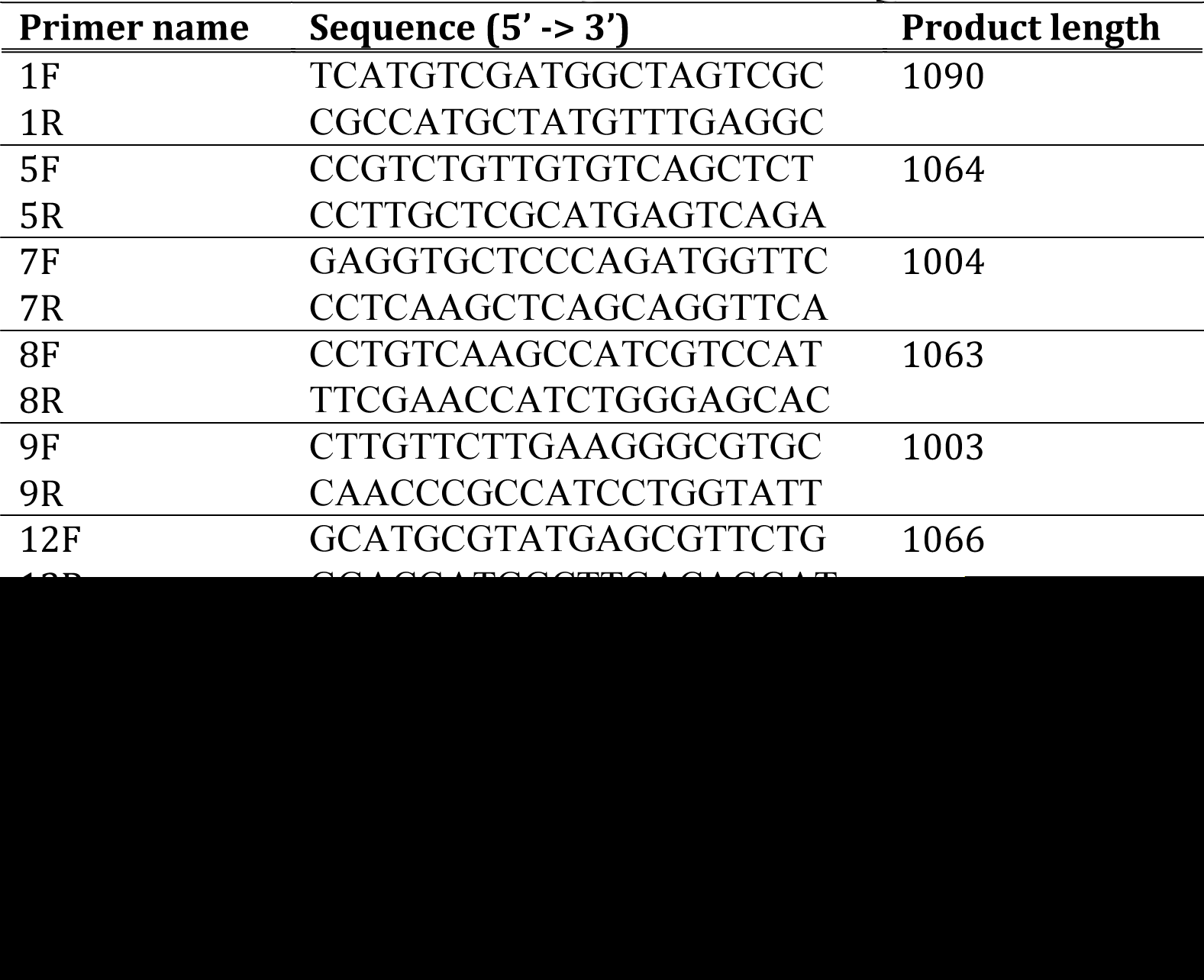
Primers used to test for the presence of MLOC_17240.

**S1 Files**. Folder containing Web archive files from Clustal Omega showing alignment of reads from both WT and MUT tissue. These were sequenced by Functional Biosciences (Madison, WI, USA) and aligned to MLOC_17240 (HORVU5Hr1G120850).

